# Mechanosensitive ion channel MSL8 is required for pulsatile growth and cell wall dynamics in *Arabidopsis* pollen tubes

**DOI:** 10.1101/2023.07.27.550874

**Authors:** Joshua H. Coomey, Elizabeth S. Haswell

## Abstract

**HIGHLIGHT:** Pollen tube growth requires tight control of apical wall expansion. We present evidence for a mechanosensitive ion channel, MSL8, as a braking signal in growth dynamics through cell wall regulation.

The male gametophyte in flowering plants, pollen, both performs the critical role of fertilization and represents a unique and accessible system for interrogating plant cell mechanics. Pollen endures multiple mechanical hurdles during its lifecycle: desiccation in the anther, rapid rehydration on the stigma, and germination to produce a rapidly growing pollen tube that will eventually reach the ovule. A key component in this robust mechanical system is MscS-Like 8 (MSL8), a mechanosensitive ion channel. We previously proposed that that MSL8 serves as an “osmotic safety valve”, regulating pressure in the germinating pollen tube by releasing anions in response to plasma membrane tension, thereby preventing pollen tube rupture. However, we subsequently identified defects in the cell walls of *msl8* mutant pollen, suggesting that it plays a role independent of osmoregulation, a conclusion also supported by mathematical modeling. Here, we show that pollen tubes lacking MSL8 channel function by genetic knockout or channel-blocking point mutation lose major growth pauses, have altered pectin esterification patterns, and are sensitive to pectin crosslinking. Together, these data suggest a mechanism whereby tension-gated ion release through mechanosensitive channels regulates apoplastic function and cell wall dynamics.

## INTRODUCTION

Pollen is the male gametophyte of flowering plants. Successful pollination is required for the production of all seed and fruit crops we reap from angiosperms, as well as for the propagation of the next generation (Johnson *et al*., 2019). During the process of pollination, pollen grains undergo a drastic mechanical journey: they develop and fully desiccate in the anther, then travel to the female pistil where they rapidly rehydrate. After rehydration, the pollen grain germinates a pollen tube, one of the fastest growing cells in the plant kingdom (Shamsudhin *et al*., 2016). The tube then travels to the ovule to deliver the sperm cells to the egg cells for fertilization. This sequence of events requires tight regulation of pollen grain and tube development and growth, which must be additionally coordinated with the perception of, and adaptation to, huge changes in osmotic conditions and substantial mechanical constraints (Williams *et al*., 2016; Cameron and Geitmann, 2018; Reimann *et al*., 2020).

Unlike most plant cells, pollen tubes are tip-growing, meaning that they elongate by cell wall deposition at the cell apex rather than across the cell length as in anisotropic expansion (Hepler *et al*., 2001; Winship *et al*., 2021). Controlled tip-growth requires a balance between increased cell volume and the delivery and incorporation of new cell wall material. Cell expansion and cell wall deposition are coordinated by numerous intracellular factors, including ion fluxes, cytoskeleton dynamics, the accumulation of reactive oxidative species, changes in turgor, and multiple signaling pathways (Johnson *et al*., 2019). Imbalance between expansion and cell wall deposition can result in a loss of cell integrity or a lack of growth (Johnson *et al*., 2019).

Pollen tubes can grow continuously or in a pulsatile manner. Their rate of growth varies widely both between and within a species, with rates ranging from 100-600 nm/s in lily (Cárdenas *et al*., 2008), 10-50 nm/s in tobacco (Michard *et al*., 2008), and 30-150 nm/s in *Arabidopsis thaliana* (Rounds *et al*., 2011; Damineli *et al*., 2017). The robustness of growth oscillations also varies: lily pollen exhibits highly periodic elongation in 20-50 second cycles (Cárdenas *et al*., 2008), while tobacco pollen has ∼80 second periods (Michard *et al*., 2008). Pollen tubes from *A. thaliana* show relatively less uniform oscillatory characteristics, with reports ranging from ∼30-90 second cycles, ambiguous oscillations, or steady growth rates (Rounds *et al*., 2011; Damineli *et al*., 2017).

The internal expansive force that drives the pollen tube tip forward has long been attributed to turgor. However, turgor does not appear to be responsible for growth oscillations. Multiple studies have suggested that turgor pressure remains constant in growing pollen tubes, even those exhibiting pulsatile behavior. This has led to the hypothesis that cyclic changes in the cell wall, either through deposition or modification of cell wall components, modulate the pulsatile growth dynamics of pollen tubes (Benkert *et al*., 1997; Zerzour *et al*., 2009; Winship *et al*., 2010, 2021; Rojas *et al*., 2011; Van Hemelryck *et al*., 2018; Dumais, 2021).

The wall composition of pollen tubes differs from most plant tissues in several respects. Callose is a significant component of pollen tube walls, where it is found along the tube shank and making up thick plugs that segment the growing pollen tube, keeping cytoplasmic space compacted towards the tip (Parre and Geitmann, 2005; Chebli *et al*., 2012). In addition, in pollen tubes the deposition, modification, and crosslinking of pectin, rather than the orientation of cellulose microfibrils, stabilizes and directs growth (Hepler *et al*., 2001; Aouar *et al*., 2010).

Pectins are a large class of diverse cell wall polysaccharides (Wolf *et al*., 2009; Harholt *et al*., 2010). The most common pectin found in pollen tubes is homogalacturonan. This ɑ-1,4-linked galacturonic acid homopolymer is synthesized in the Golgi with a high degree of methyl-esterification. The degree of homogalacturonan esterification directly impacts the extent to which that homogalacturonan polymer is integrated and crosslinked with other polymers in the cell wall. The highly esterified homogalacturonan delivered in vesicles to the tube apex has a low capacity for crosslinking, making a more extensible wall (Giovane *et al*., 2004; Bosch and Hepler, 2005; Guo *et al*., 2022).

Pectin-modifying proteins, such as pectin methylesterases (PMEs) and pectin methylesterase inhibitors (PMEIs), are co-delivered in vesicles with esterified homogalacturonan. When active, PMEs cleave the methyl group of homogalacturonan, leaving negatively charged, de-esterified homogalacturonan. Calcium ions in the wall space complex with these negatively charged pectins, crosslinking them into an “egg box” configuration, and thereby stiffening the pectic gel matrix, and making the wall less extensible (Bosch *et al*., 2005; Bosch and Hepler, 2006; Leroux *et al*., 2015).

This enzymatic conversion from esterified homogalacturonan to de-esterified, cross-linkable homogalacturonan is spatially regulated to maintain an extensible tube tip and rigidified tube shank. While PMEs appear to localize throughout the pollen tube wall, their activity is tightly regulated, with lower activity at the tube apex (Bosch *et al*., 2005; Röckel *et al*., 2008). For example, changes in cell wall microenvironment, particularly pH and ionic strength, can cause PMEs to conjugate with the cell wall or PMEIs and restrict their catalytic activity (Bosch and Hepler, 2005; Bonavita *et al*., 2016).

Mechanosensitive (MS) ion channels also play an important role in pollen. It is well-established that, by opening a pore in response to lateral membrane tension, MS channels can function as osmotic regulators (by allowing ions and osmolytes to exit the cell) and as signal transducers (by allowing calcium to enter the cell) (Peyronnet *et al*., 2014). Two *Arabidopsis* MS ion channels, MscS-Like 8 (MSL8) and close homolog MSL7, are expressed in pollen grains and pollen tubes. Loss of MSL8 results in increased rates of pollen grain germination and bursting, with altered callose deposition in germinating grains (Hamilton *et al*., 2015; Wang *et al*., 2022). A *bona fide* mechanosensitive channel, MSL8 shows a slight preference for anions (Hamilton *et al*., 2015; Hamilton and Haswell, 2017). Single amino acid mutations are sufficient to disrupt MSL8 function, particularly F720L, which blocks channel conductance (Hamilton and Haswell, 2017). MSL8^F720L^ mutants showed similar phenotypes as a *msl8* knockout, indicating that channel activity is required for proper function (Hamilton *et al*., 2015; Hamilton and Haswell, 2017). No phenotypes have yet been shown in plants lacking MSL7.

While osmoregulation is a core property of MscS-like proteins across systems (Hurst *et al*., 2008), and we previously proposed that MSL8 may function as an osmotic safety valve (Hamilton *et al*., 2015), recent evidence suggests that MSL8 may play a role beyond or independent of simple osmoregulation. Germinating *msl8* mutant pollen grains have altered cell wall deposition, with increased callose at the germination site and a non-uniform distribution of cell wall material in the germination plaque (Wang *et al*., 2022). Overexpressing MSL8 leads to excess callose deposition in the periphery of ungerminated grains, possibly as part of a genetic interaction between *MSL8* and genes that encode components of the cell wall integrity (CWI) pathway (Wang *et al*., 2022).

Finally, mathematical modeling of pollen germination also suggested that MSL8 may have functions in addition to or instead of osmoregulation. During rehydration, *msl8* mutant pollen grains swell and deform to a greater extent than wildtype, which plateaus in volume (Miller *et al*., 2022). Mathematical modeling can simulate this process with contributions of various factors, such as MSL8 tension-gated osmoregulation, turgor, and cell wall deformation. Kinetic models best support a scenario where cell wall modification leads to the observed changes in hydration dynamics, rather than simple osmoregulation (Miller *et al*., 2022).

Here, we build on these data to investigate the relationship between MSL8 ion channel function, pollen tube growth, and cell wall composition. These experiments serve to further test the hypothesis that MS ion channels are capable of altering cell wall extensibility as well as osmoregulation and intracellular signaling.

## METHODS

### Plant growth

All lines were in the Col-0 *Arabidopsis thaliana* ecotype and validated in previous publications (Hamilton *et al*., 2015; Hamilton and Haswell, 2017; Wang *et al*., 2022; Miller *et al*., 2022). Seeds were sown directly on hydrated Berger BM2 growth medium and stratified for 24 hours at 4°C. Flats were then moved to a Conviron MTPS120 growth chamber and grown in long day conditions (16h light, 8h dark) at 21°C, 175 µm light, and 50% relative humidity. Plants began to bolt ∼5 weeks after sowing, with pollen collected 5-10 days after bolting.

### In vitro pollen germination

For each genotype, 20-40 freshly opened flowers were collected into 1.7 ml Eppendorf tubes containing 750 µl of liquid pollen growth medium (LPGM, 18% sucrose, 2 mM CaCl_2_, 2 mM Ca(NO_3_) _2_, 0.49 mM H_3_BO_3_, 1 mM MgSO_4_, 1 mM KCl, pH’d to 7.05 with KOH (Bou Daher *et al*., 2009; Wang *et al*., 2022)). Tubes were vortexed at maximum speed for 1 min to release pollen, then centrifuged at 10,000xg for 4 min to pellet pollen. Forceps were used to remove excess floral debris and the supernatant was carefully removed by pipette.

For germination and growth in Eppendorf tubes, the pollen pellet was resuspended in 200 µl LPGM and tubes transferred to a petri dish humidified with a wet Kimwipe. For germination and growth on microwell dishes, coverslip bottom microwell dishes (MatTek Corp, no. P35G-1.5-14-C) were coated with 150 µl of poly(ethyleneimine) solution (1:30 in ddH_2_O) for 10 min, washed twice with 200 µl ddH_2_O, and coated with 200 µl of LPGM. Following the pollen centrifugation described above, the pollen pellet was resuspended in 20 µl LPGM. The 200 µl LPGM was removed from the dish and the resuspended pollen added with a cut pipette tip. The pollen was left for 1 min to adhere to the plate, then 200 µl fresh LPGM was added carefully by pipette. Microwell dishes were placed in petri dishes humidified with a wet Kimwipe in an incubator at 23°C.

### Pollen tube growth dynamic assays and analysis

Pollen was germinated in microwell dishes as described above and grown for 4 hours before imaging on an Olympus IX73 microscope equipped with a Hamamatsu C11440 digital camera. Brightfield images were captured with a 10x / NA0.3 objective at 10 second intervals over a 40-minute period. Kymograph analysis was carried out in FIJI, using a segmented line with width = 1 drawn down the center longitudinal axis of the pollen tube, and processed with the Multi Kymograph function. Kymographs were analyzed using the CHUKKNORRIS R scripts (Damineli *et al*., 2017). To convert kymographs to a timeseries, images were thresholded in ImageJ such that the tip-boundary was defined by black pixels. All pixels below the tip-boundary were manually converted to black and all pixels above were manually converted to white. The kymograph was oriented such that time increased down the Y axis and tube length increased along the X axis. These images were saved as .txt files, with black pixels = 255 and white = 0. These files were read into R and converted to a data frame of time and length by assigning time values to columns and summing pixel values across rows for tube length. This data frame could then be loaded into the CHUKKNORRIS AnalyzeTimeSeries script, with the smooth.tm value adjusted to 30. CHUKKNORRIS output tables were used to calculate average maximum and minimum rates.

### Pharmacology

For PMEI treatment, epigallocatechin gallate (EGCG, (−)-Epigallocatechin gallate ≥ 95%, Sigma-Aldrich E4143) from green tea was used as a pectin methylesterase inhibitor (PMEI) (Lewis *et al*., 2008). The ECGC was resuspended in water at a 100 mM stock concentration. This was added to LPGM for a final concentration of 0.1 mM. LPGM adjusted with an equal volume of water was used as the mock treatment.

For PME treatment, pectinesterase from orange peel (Sigma-Aldrich P5400) was resuspended in 1 M phosphate buffer, pH 7.5 at 1000 U/ml. This was added to LPGM for a final concentration of 1 U/µl. LPGM adjusted with an equal volume of 1 M phosphate buffer was used as the mock treatment.

Pollen was germinated in microwell dishes as described above and grown for 2 hours before pharmacological treatment. Then LPGM was removed from the dish by gentle pipetting and replaced with 200 µl of treatment or mock solution as described above. Treated pollen was grown for an additional hour, then imaged and analyzed in the kymograph assay described above.

### Immunolabeling

Pollen was germinated and grown in 1.7 ml Eppendorf tubes as described above for 5 hours. Monoclonal antibody labeling of esterified and de-esterified homogalacturonan was carried out based on previously described protocols with the minor modifications detailed here (Mecchia *et al*., 2017). Tubes were then centrifuged at 4000xg for 5 minutes to collect the pollen, and LPGM removed by pipette. This combination of centrifugation and pipetting is shared for all subsequent solution exchange steps unless noted otherwise. Pollen was fixed by resuspending gently in 200 µl PEM buffer (4% paraformaldehyde in 1 M NaOH, 50 mM PIPES, 1 mM EGTA, 5 mM MgSO_4_, pH6.9) at room temperature overnight. The fixative was removed, and the pollen washed with 1x PBS (137 mM NaCl, 10 mM Na2HPO4.2H_2_0, 1.8 mM KH2PO4, 2.7 mM KCl, pH 7.4). The pollen was then incubated in 1x PBS + 5% milk protein (1x PBS+MP) for 1 hour at room temperature, then incubated with a 1:100 dilution of LM20 or LM19 primary antibody (Kerafast, ELD003 and ELD001 respectively) (Verhertbruggen *et al*., 2009; Yang *et al*., 2022) in 1x PBS+MP for 1 hour at room temperature. The pollen was then washed 3 times in 1x PBS and incubated in a 1:1000 dilution of rabbit anti-rat IgG FITC (Sigma, F1763-.5ML) in 1x PBS+MP for 1 hour. The pollen was washed 3 times with 1x PBS and then mounted on glass slides for imaging. Images were captured on an Olympus Fluoview3000 point scanning confocal laser microscope with a 60x / NA1.2 water objective and HyD detectors, using excitation at 488 nm and emission collection at 510-530 nm. Intensity profiles were analyzed in ImageJ by tracing a meridional line along the periphery of the cell wall starting at the tube apex and proceeding at least 50 µm down the shank.

### Toluidine Blue staining

Pollen was germinated in the microwell dish setup described above and grown for 4 hours. LPGM was replaced with LPGM + 0.01% w/v Toluidine Blue (Sigma-Aldrich 198161-5G) and incubated for 20 minutes. Pollen tubes were then imaged on an Olympus IX73 microscope in brightfield with a 10x / NA0.3 objective and Hamamatsu C11440 digital camera.

### Aniline blue staining

Pollen was germinated in the Eppendorf tube setup described above and grown for 4 hours. LPGM was replaced with acetic acid + 10% ethanol and fixed for 2 h. The fixative was replaced with 1M NaOH and incubated at room temperature overnight. Following two washes in KPO buffer (4mM K_2_HPO_4_, 0.83mM KH_2_PO_4_, pH 7.5) the fixed pollen tubes were stained with 0.01% decolorized aniline blue for 3 h. Images were captured on an Olympus Fluoview3000 point scanning confocal laser microscope with a 60x / NA1.2 water objective and HyD detectors, using excitation at 405 nm and emission collection at 430-470 nm. Intensity profiles were analyzed in ImageJ by tracing a meridional line along the periphery of the cell wall starting at the tube apex and proceeding at least 50 µm down the shank.

### Direct Red 23 staining

Pollen was germinated in the microwell dish setup described above and grown for 4 hours. LPGM was replaced with Direct Red 23 (10µg/ml) in 1x PBS and incubated for 5 min. Images were captured on an Olympus Fluoview3000 point scanning confocal laser microscope with a 60x / NA1.2 water objective and HyD detectors, using excitation at 561nm and emission collection at 580-615 nm. Intensity profiles were analyzed in ImageJ by tracing a meridional line along the periphery of the cell wall starting at the tube apex and proceeding at least 50 µm down the shank.

### Cos-488 staining

Pollen was germinated in the microwell dish setup described above and grown for 4 hours. LPGM was replaced with LPGM + Cos-488 (1:300) and incubated for 5 min. Images were captured on an Olympus Fluoview3000 point scanning confocal laser microscope with a 60x / NA1.2 water objective and HyD detectors, using excitation at 488nm and emission collection at 510-530 nm. Intensity profiles were analyzed in ImageJ by tracing a meridional line along the periphery of the cell wall starting at the tube apex and proceeding at least 50 µm down the shank.

### Statistical analysis

Data analysis and statistical testing was carried out using R (version 4.1.1) (Wickham and Grolemund, 2017), with packages ‘ggplot2’ used for data visualization, ‘dplyr’ and ‘data.table’ used for data transformation, and ‘multcompView’ for generating compact letter displays.

## RESULTS

### MSL8 channel function is required for pulsatile growth dynamics

To explore the effects of MSL8 channel function on pollen tube growth, we leveraged a suite of genetic reagents including functional *MSL8* (WT), a *msl8* CRISPR knockout mutant (*msl8-5, (Wang et al., 2022)*), a *msl8* mutant complemented with natively expressed functional MSL8 (*msl8-5 + pMSL8::MSL8, (Miller et al., 2022)*) and a *msl8* mutant complemented with natively expressed non-functional MSL8 (*msl8-5 + pMSL8::MSL8^F720L^*,(Hamilton and Haswell, 2017; Miller *et al*., 2022)) (Figure 1A, left panels). Pollen tubes were germinated and grown in microwell dishes for 3 hours before time-lapse imaging every 10 s for 40 min. From each time-lapse series we created kymographs of individual tubes elongating (Figure 1A, right panels).

**Figure 1.**
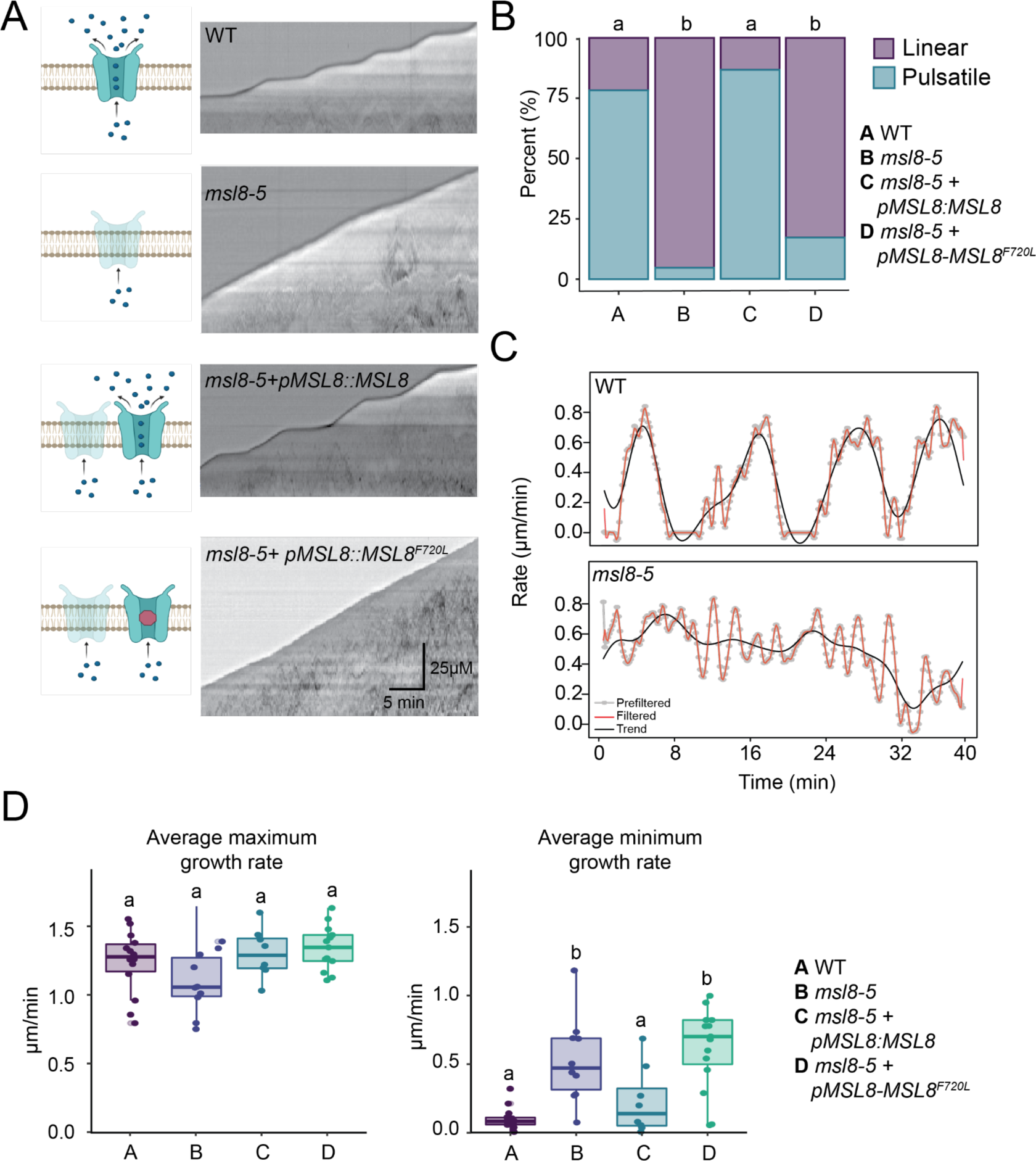
Pollen tubes lacking MSL8 channel function lose pulsatile growth dynamics. **A)** Left panels, schematic representation of MSL8 variant channel functions. Right panels, kymograph analysis of pollen tube elongation over 40 min of growth. Tubes were imaged every 10 s. **B)** Quantification of linear vs pulsate growth dynamics in pollen tubes with or without functional MSL8, n=30-40 tubes per genotype. Statistical analysis by Fisher’s Exact test and represented by compact letter display, p<0.05. **C)** Representative traces of growth rates extracted from kymograph analysis, top panel showing pulsatile growth and bottom panel showing linear growth. **D)** Average maximum (left) and minimum (right) instantaneous growth rates, derived from kymograph analysis, n=12-22 tubes per genotype. Statistical analysis by ANOVA followed by Tukey HSD testing and represented by compact letter display, p<0.05.

We observed wildtype *A. thaliana* pollen tubes with both pulsatile and linear elongation patterns, with most tubes (∼75%) having some degree of pulsatile behavior and a minority (∼25%) elongating continuously **(Figure 1B-C, Supplemental Figure 1)**. In *msl8-5* mutants, however, nearly all pollen tubes elongated continuously without major growth pauses **(Figure 1B-C, Supplemental Figure 1)**. We observed this loss of pulsatile growth in two other *msl8* mutant lines, while single *msl7* mutants retained pulsatile growth patterns similar to wildtype **(Supplemental Figure 1)**. Functional complementation of *msl8-5* with *pMSL8::MSL8* restored pulsatile growth in three independent lines, while three independent lines of the channel-blocked *MSL8^F720L^* failed to restore pulsatile growth to *msl8-5* mutant pollen tubes (**Figure 1B-C, Supplemental Figure 1)**.

To quantify changes in growth rate, we further analyzed our kymograph results using the CHUKKNORRIS R scripts (Damineli *et al*., 2017). Representative growth rate traces of wildtype and *msl8-5* pollen tubes are depicted in **Figure 1C**, upper and lower panels respectively. The average instantaneous maximum growth rate across all genotypes was not significantly different between genotypes **(Figure 1D**, left panel). However, pollen tubes lacking MSL8 channel function (both *msl8-5* and *msl8-5 + pMSL8::MSL8^F720L^*) had a higher average instantaneous minimum rate than wildtype or functionally complemented lines **(Figure 1F**, right panel). These data provide an explanation for our previous observation that *msl8* pollen tubes grow longer than wildtype, which we previously interpreted as resulting from a higher growth rate (Hamilton *et al*., 2015; Wang *et al*., 2022).

To summarize, tubes lacking MSL8 channel function had similar maximum instantaneous growth rates as tubes with functional MSL8, but lacked major growth pauses, suggesting a role for MSL8 in promoting elongation arrest. Taken together, these data indicate that MSL8 channel function is required for pulsatile growth dynamics in in vitro germinated pollen tubes.

### Osmotic support does not suppress the loss of growth pauses in *msl8* mutants

One way in which a MS ion channel could be required for pollen tube pausing could be through an effect on osmoregulation, an established function of MscS-like channels (Levina *et al*., 1999; Flegler *et al*., 2022). As a mechanosensitive anion efflux channel, MSL8 may serve to relieve internal osmotic pressure as the pollen grain rehydrates and swells. Indeed, we previously showed that osmotic support suppresses the *msl8* mutant phenotype of increased grain bursting during hydration (Hamilton *et al*., 2015). Similarly, the continuous elongation of pollen tubes lacking MSL8 channel function observed in Figure 1 could be the result of defective osmoregulation. To test this possibility, we grew pollen tubes in LPGM supplemented with 0, 50 or 100 mM mannitol. Kymograph analysis showed that *msl8-5* pollen did not show additional growth pauses with osmotic support **(Figure 2A)**. However, osmotic support did reduce average maximum growth rate and average minimum growth rate in both wildtype and *msl8-5* pollen **(Figure 2B)**. Thus, while increased osmolarity of the growth medium did reduce elongation rate generally, it did not affect the lack of pausing in *msl8* mutants.

**Figure 2.**
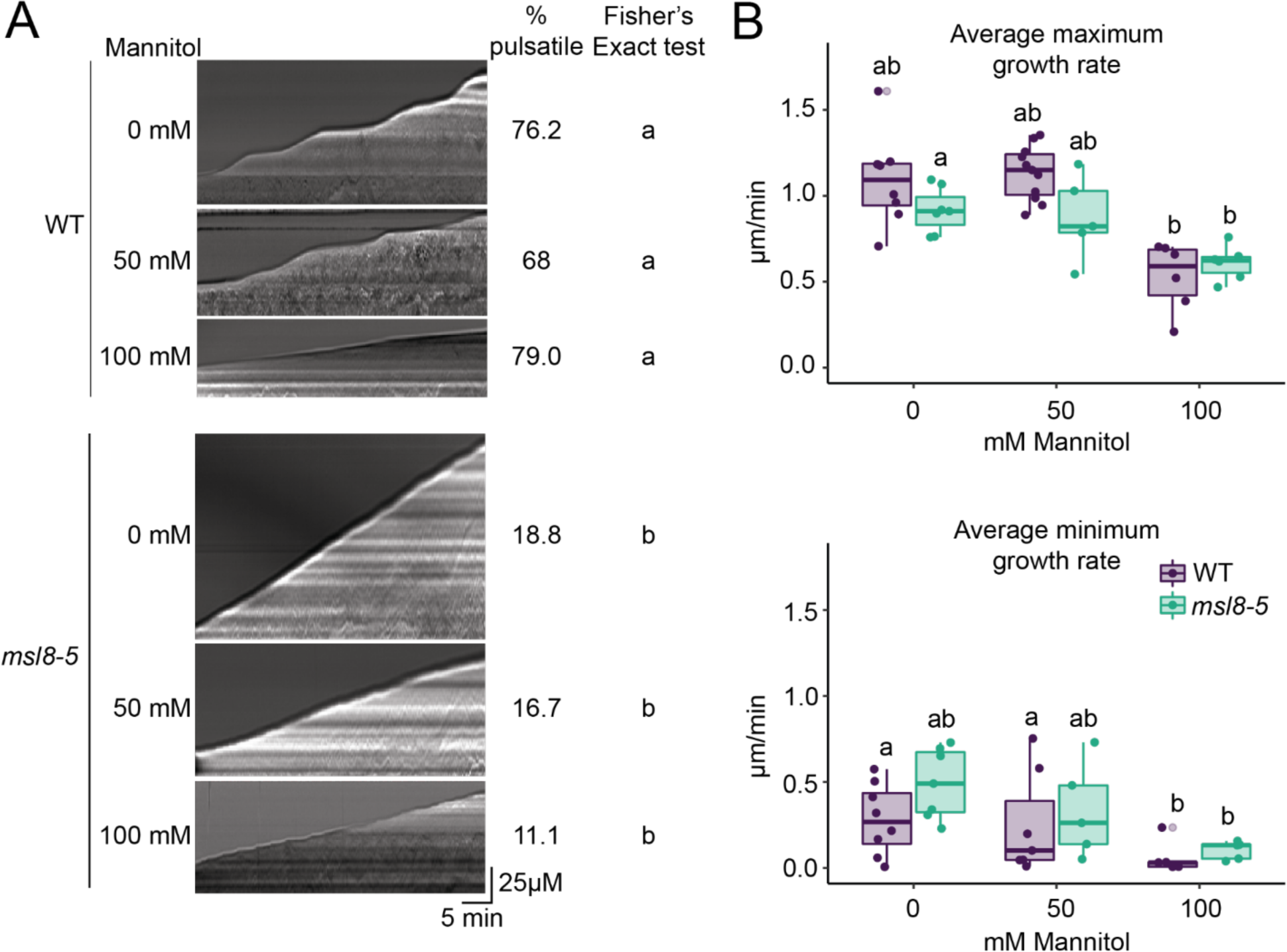
Osmotic support does not restore pulsatile growth dynamics in *msl8* pollen tubes. **A)** Kymograph analysis of pollen tube elongation over 40 min of growth in media with increasing concentrations of mannitol. Tubes were imaged every 10 s. The percent of tubes with pulsatile growth was calculated and listed tithe right of representative images. Statistical analysis by Fisher’s exact test, with significance denoted by compact letter display; p<0.05. **B)** Average maximum (left panel) and minimum (right panel) growth rate from kymograph analysis, n=20-25 tubes per genotype. Statistical analysis by ANOVA followed by Tukey HSD testing and represented by compact letter display, p<0.05.

### MSL8 channel function is required for normal cell wall patterning

We next considered that the lack of growth pauses in *msl8* mutant pollen tubes might result from defects in dynamic cell wall modifications associated with oscillatory growth. Pollen tube growth pauses are associated with the accumulation of new cell wall material, which produces a banding pattern when stained with the general cell wall dye toluidine blue (Hoedemaekers *et al*., 2015).

We germinated and grew pollen tubes in microwell dishes for 4 hours before staining with toluidine blue and imaging in brightfield with a color camera. We observed the expected banded pattern in WT and *msl8-5 + pMSL8::MSL8* pollen tubes, while in tubes from *msl8-5* mutants or *msl8-5 + pMSL8::MSL8^F720L^* mutants we observed a smooth staining pattern lacking notable bands of cell wall deposition **(Figure 3)**. These banded and smooth cell wall patterns were observed at frequencies similar to the pulsatile and linear elongation patterns seen in our kymograph analysis **(Figure 1B)**, and held true for multiple *msl8* mutant alleles, and multiple independent transgenic lines **(Supplemental Figure 2)**. Thus, MSL8 channel function is required for the typical cell wall patterning associated with pulsatile growth.

**Figure 3.**
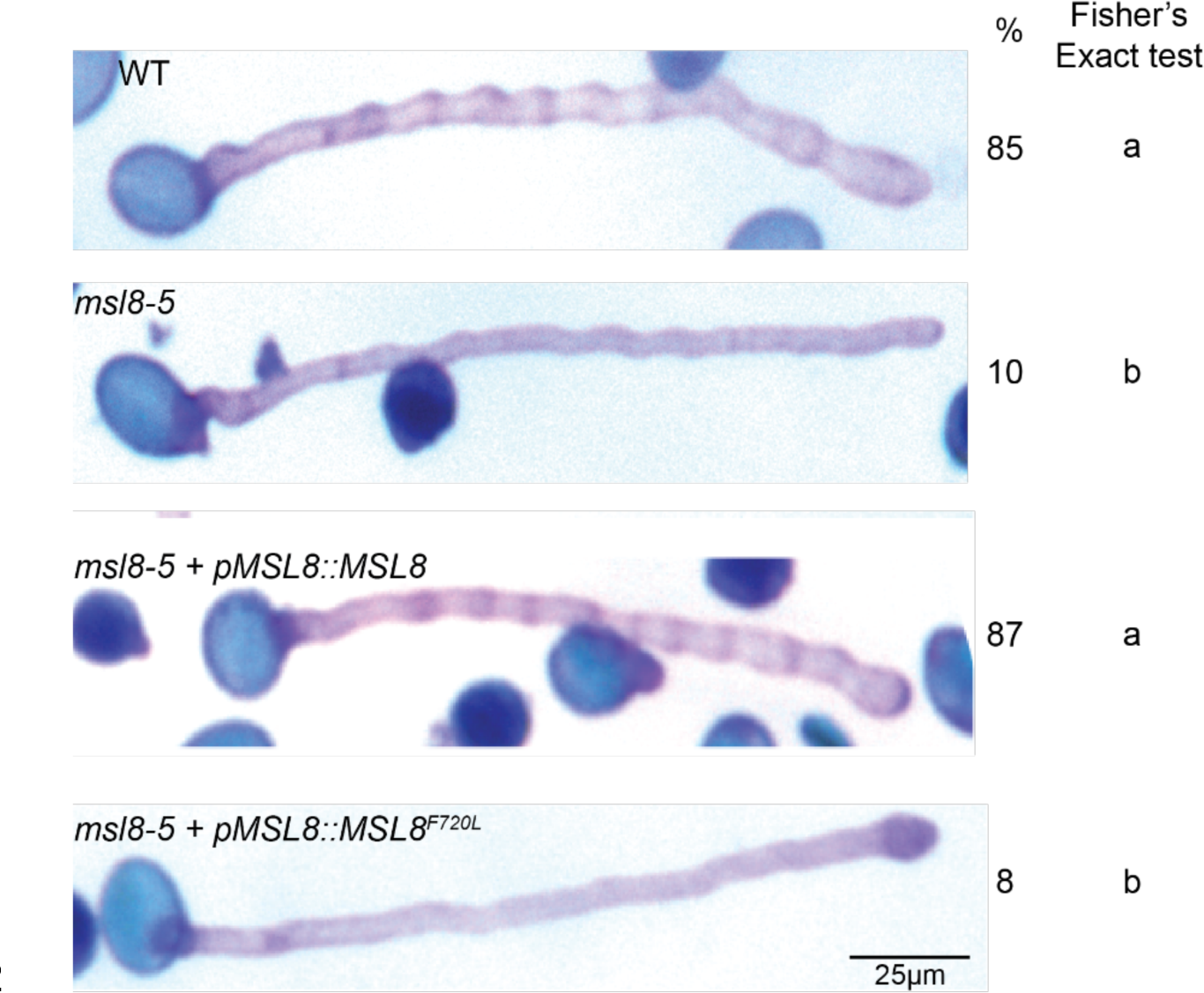
Pollen tubes lacking MSL8 channel function lack the distinctive WT banding pattern when stained with toluidine blue. Images of toluidine blue-stained pollen tubes. On the right of each image is the % of pollen tubes that showed banding for each genotype, n = 100 tubes per genotype. Statistical analysis by Fisher’s exact test, with significance denoted by compact letter display; p<0.05.

### Callose and cellulose deposition are unchanged in *msl8* pollen tubes compared to the WT

Given our previous observation of altered callose accumulation in *msl8* pollen grains (Wang *et al*., 2022), we next investigated callose and cellulose levels in *msl8* pollen tubes. To detect callose levels, we grew pollen tubes for four hours before fixing and staining with decolorized aniline blue and imaging with confocal microscopy (Wang *et al*., 2022). Only a slight increase in shank callose levels in *msl8-5* pollen tubes was observed compared to the WT **(Figure 4A)**. To detect cellulose levels, we grew pollen tubes for four hours before fixing and staining with Direct Red 23 (also known as Pontamine Fast Scarlet 4B (Anderson *et al*., 2010; Bidhendi *et al*., 2020)). No notable difference in cellulose levels or patterning between the genotypes was observed **(Figure 4B)**.

**Figure 4.**
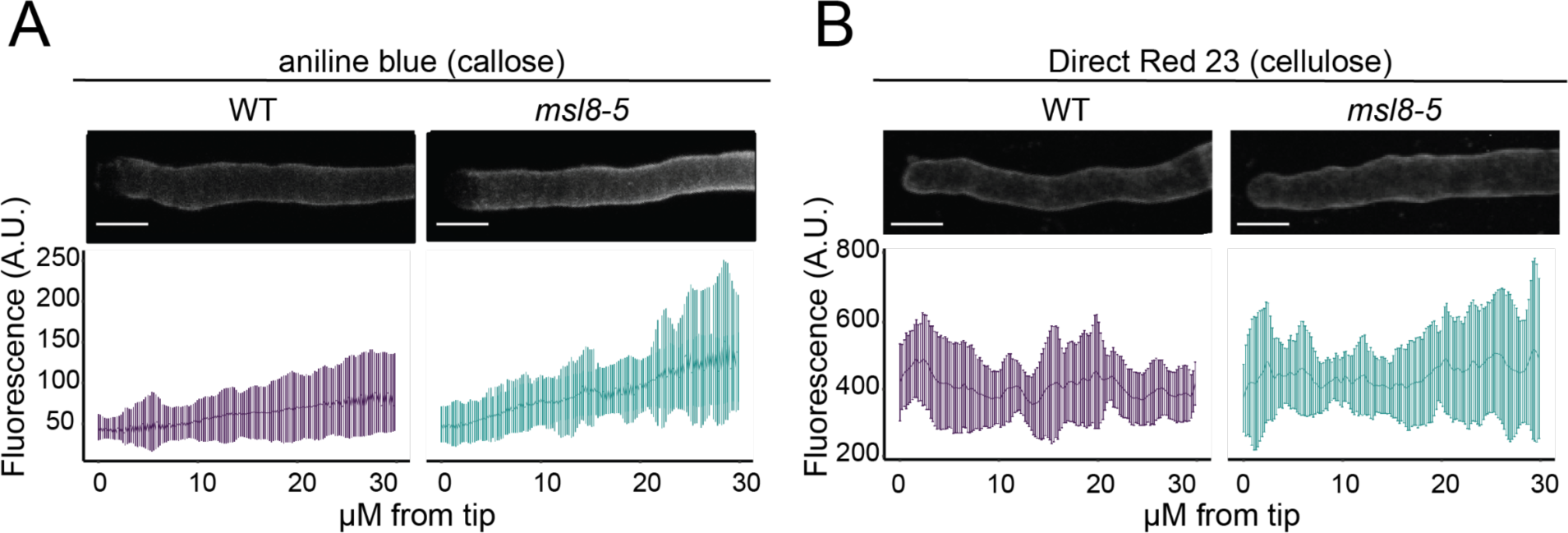
Pollen tubes lacking MSL8 channel function have slightly increased shank callose levels and similar cellulose levels compared to wildtype. **A)** Top, representative maximum-projection images of aniline blue staining in pollen tubes fixed after 7h of growth. Bottom, quantification of aniline blue signal (arbitrary units, A.U.) along the meridian of the pollen tube, measured from the apex along the cell wall for 30 microns. The trend line represents the mean fluorescent signal, with error bars representing standard deviation. **B)** Top, representative maximum-projection images of Direct Red 23 staining of cellulose in pollen tubes fixed after 7h growth. Bottom, quantification of Direct Red 23 signal (arbitrary units, A.U.) along the meridian of the pollen tube, measured from the apex along the cell wall for 30 microns. The trend line represents the mean fluorescent signal, with error bars representing standard deviation. n=40 tubes per genotype. Scale bar, 10 μm.

### MSL8 channel function is required for wild-type pectin esterification patterning

The deposition, de-esterification, and crosslinking of pectin are key steps in pollen tube tip growth. Having observed that tubes lacking MSL8 channel function have altered growth dynamics and changed cell wall patterning, we next asked if pectin patterning may be impacted in these lines.

Esterified homogalacturonan, considered more extensible due its lack of crosslinking capacity, is deposited at the tube apex and can be detected by LM20 antibody (Verhertbruggen *et al*., 2009; McKenna *et al*., 2009; Chebli *et al*., 2012; Bidhendi *et al*., 2020). We grew pollen tubes for 5 hours in Eppendorf tubes before fixing and immunostaining with LM20 primary antibody and a fluorescently tagged secondary antibody. We then measured the fluorescent signal by confocal microscopy and quantified the signal intensity along the tube wall from the apex to 20 µM down the shank. In WT tubes, we observed the expected pattern of esterified homogalacturonan restricted to the tube apex. However, tubes lacking MSL8 channel function had more intense LM20 antibody signal at the tube apex and further down from the tube tip than the WT, indicating higher esterified homogalacturonan levels over a larger region of the tube. This pattern was also observed in channel-blocked complementation lines. Functionally complemented lines had slightly less LM20 signal at the tube apex than WT, possibly due to mild overexpression **(Figure 5A, Supplemental Figure 3A)**.

**Figure 5.**
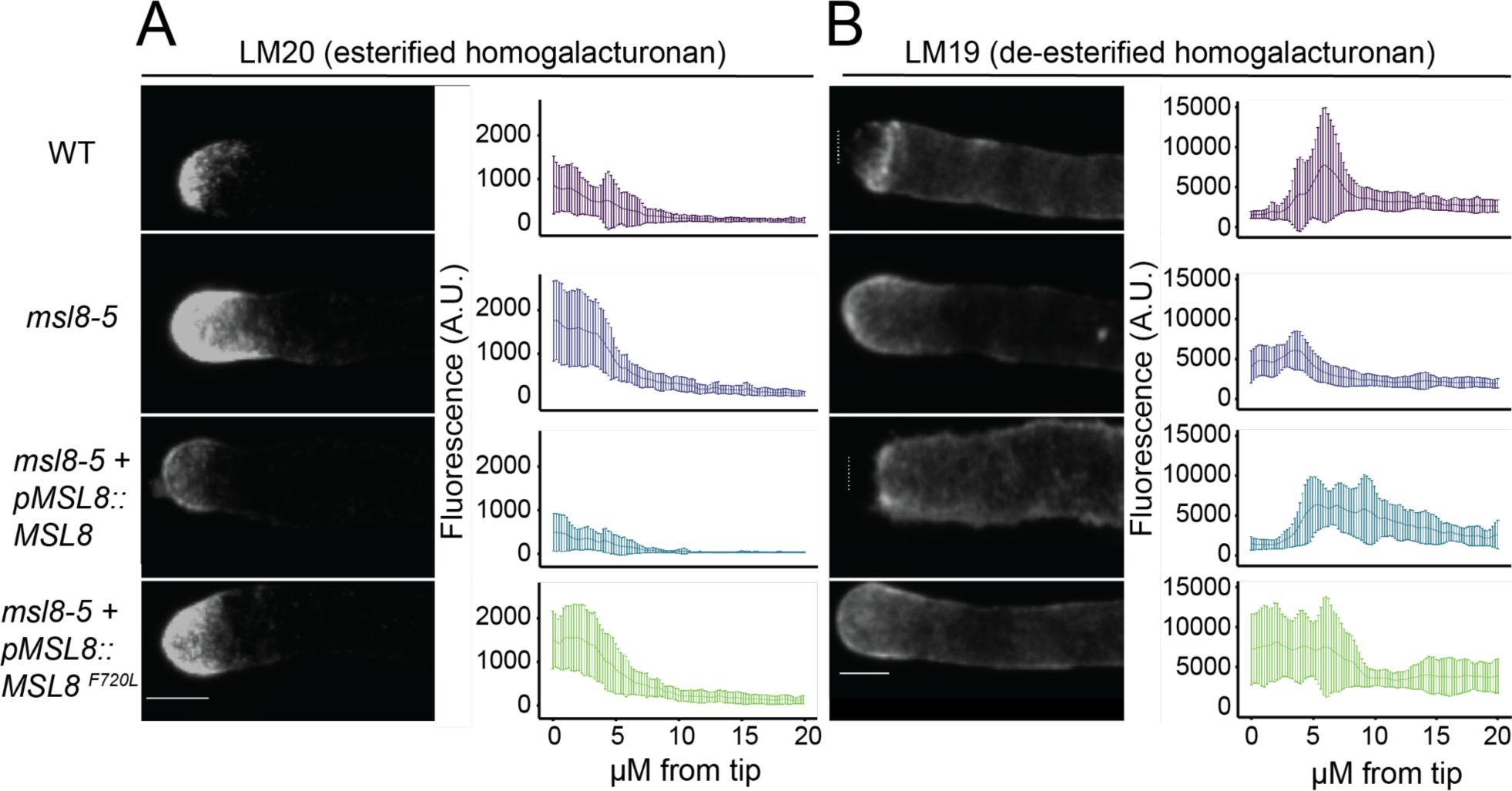
Pollen tubes lacking MSL8 channel function have altered pectin distribution. Representative maximum-projection images of immunolabeled pectins. Left panels, LM20 antibody labeling of esterified homogalacturonan (**A**) and LM19 antibody labeling of de-esterified homogalacturonan (**B**) in pollen tubes from the indicated genotypes. Right panels, quantification of LM19 or LM20 signal along the meridian of each pollen tube, measured from the apex along the cell wall for 20 microns. Dashed lines indicate tube apex. The trend line represents the mean fluorescent signal, with error bars representing standard deviation. n = 30-40 tubes per genotype. Size bar, 5 μm.

De-esterified homogalacturonans are typically detected along the tube shank and not in the tube apex, the inverse pattern of esterified homogalacturonan (Chebli *et al*., 2012). De-esterified homogalacturonan is considered less extensible due to increased crosslinking capacity through Ca^2+^ salt bridges (McKenna *et al*., 2009; Chebli *et al*., 2012; Bidhendi *et al*., 2020). These pectins can be detected by the LM19 antibody (Verhertbruggen *et al*., 2009). Pollen tubes were processed as described above using the LM19 primary antibody to detect de-esterified homogalacturonan. The expected pattern for de-esterified homogalacturonan was observed for both wildtype and functionally complemented lines, with signal absent in the tube apex and present along the shank. However, *msl8-5* mutants as well as the *msl8 + pMSL8::MSL8^F720L^*channel blocked lines showed LM19 signal across the tube apex **(Figure 5B, Supplemental Figure 4)**.

We also assessed pectin deposition in live tubes using Cos-488, a chitosan oligosaccharide (Cos) molecule conjugated with the Alexa-488 fluorophore that binds to de-esterified homogalacturonan in an ion-dependent manner. Positive charges on the Cos probe interact with negative charges on the de-esterified homogalacturonan with a preference for homogalacturonan with a low degree of esterification (Mravec *et al*., 2014, 2017). Bidhendi et al 2020 showed Cos-488 signal across the pollen tube apex with increasing intensity up the shank, consistent with increased de-esterification along this gradient (Bidhendi *et al*., 2020). These results replicated the loss of banding in *msl8* mutant tubes seen in Toluidine blue-stained tubes **(Supplemental Figure 4)**. However, we did not observe a notable difference in intensity between WT and *msl8* mutant tubes stained with Cos-488, nor did we see the appearance of signal at the tube apex as seen with LM19 antibody. These differences likely reflect differences in the precise aspect of pectin modification that is recognized by Cos-488 and the LM19 and LM20 antibodies.

### *msl8* mutant pollen tubes are more sensitive to propidium iodide than wildtype

The presence of LM19 signal across the tube apex in *msl8* tubes **(Figure 2C-D)** suggests de-esterified homogalacturonan in that region. However, the association of de-esterified homogalacturonan with a stiffened cell wall would prompt us to expect restricted growth once crosslinked (Rojas *et al*., 2011; Hepler *et al*., 2013). However, we did not observe slower growth in *msl8* pollen tubes, leading us to hypothesize that the homogalacturonan in *msl8* tube apices is de-esterified, but not crosslinked.

To test this, we took advantage of the competitive binding of propidium iodide (PI) with sites usually occupied by calcium ions. PI will compete for Ca^2+^ binding sites in growing pollen tubes, and this dose dependent competition has been shown to inhibit pollen tube elongation (Rounds *et al*., 2011). After three hours of growth in standard LPGM, WT and *msl8-5* mutant tubes were imaged. Then the LPGM was exchanged for LPGM supplemented with 0, 40, 60 or 80 µm PI, and tubes were grown for four more hours and imaged again. The tube length before and after treatment was measured. As shown in **Figure 6A**, there was no difference in tube length in wildtype or *msl8-5* at the pretreatment time point after 3 h of growth in LPGM (left panel). After 4 h treatment with increasing concentrations of PI, wildtype also showed no difference in tube length across treatments, while *msl8-5* tubes were longer in mock conditions but the same as wildtype under increasing PI treatment (**Figure 6A**, right panel). When the change in growth rate was normalized to the mock treatment, WT pollen tubes remained unaffected by PI treatment or showed a slight tendency towards increased growth, while at all concentrations of PI *msl8-5* tubes showed a 50-60% reduction in growth **(Figure 6B)**.

**Figure 6.**
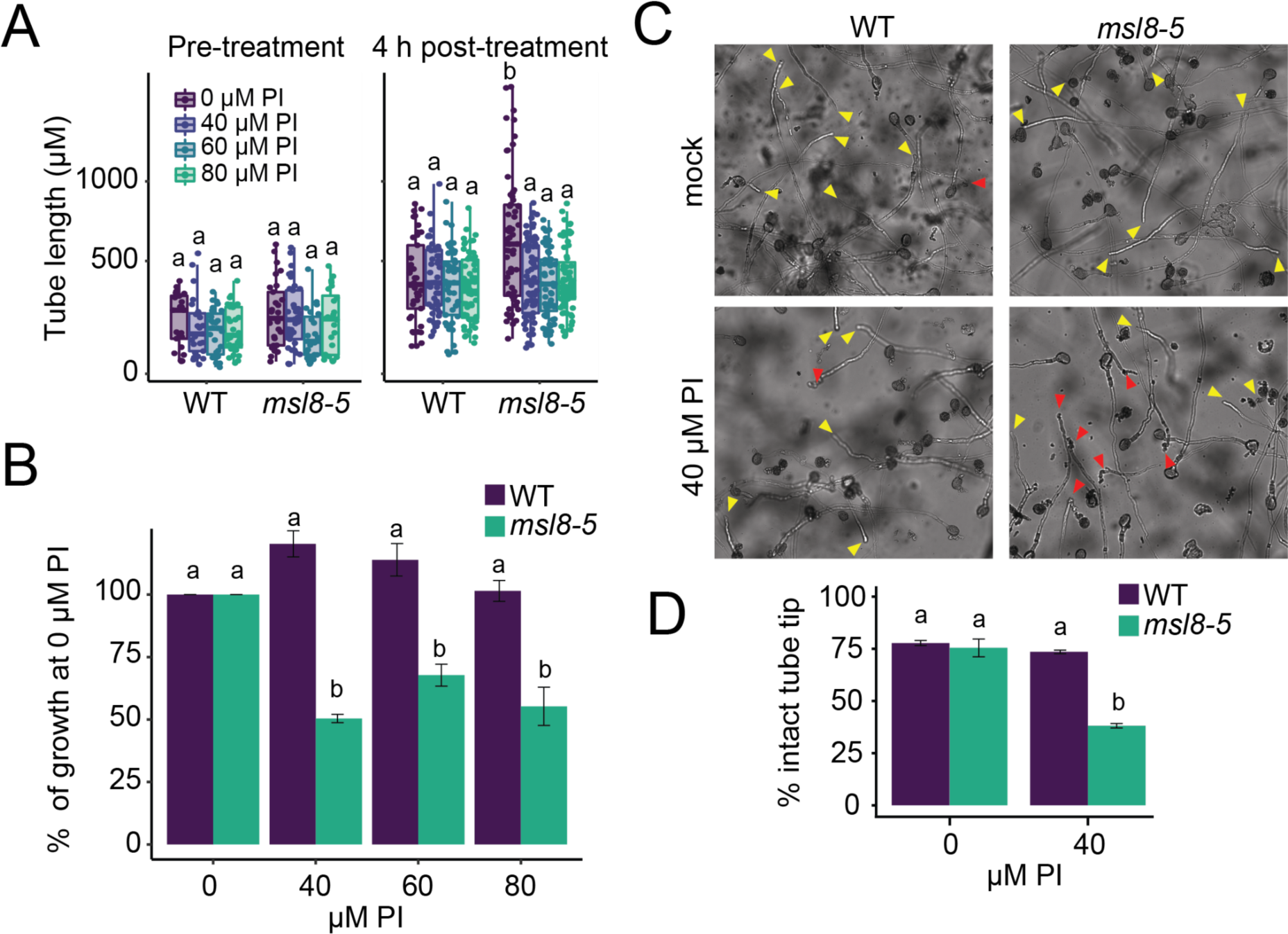
*msl8* pollen tubes are sensitive to propidium iodide (PI) treatment. **A)** Pollen tube length for wildtype and msl8-5 pollen growth pre-treatment (left) and 4 hours post-treatment (right) when grown in increasing concentrations of PI. **B)** Change in tube growth between pre- and post-treatment normalized to the control (0 µM) treatment. **C)** Reduced tube length in msl8 is associated with increased rates of tube bursting. Representative images of wildtype (left column) and *msl8-5* (right column) pollen tubes after 4 hours growth in indicated treatment. Yellow arrows indicate intact tube tips, red arrows indicate burst tube tips. **D)** Quantification of intact and tube tips in panel C. Statistical analysis by ANOVA followed by Tukey HSD testing and represented by compact letter display, p<0.05. n=40-60 tubes per genotype.

This reduction in tube length could result from growth arrest or pollen tube rupture. We quantified the number of pollen tubes with intact or burst tube tips following treatment with LPGM or LPGM supplemented with 40 µM PI and observed no difference in tube tip bursting in wildtype, but an increase in burst *msl8-5* tubes **(Figure 6C-D)**. Thus, *msl8* pollen tubes are sensitive to treatment with PI, possibly due to competition between PI and Ca^2+^ for pectin crosslinking.

### Pharmacological manipulation of PME/PMEI activity impacts WT and *msl8* growth dynamics

We next explored the effect of PME/PMEI function on the *msl8* growth phenotype. Given the broad nature of PME/PMEI function across many genes expressed in *Arabidopsis* pollen tubes, we chose a pharmacological rather than genetic approach. Epigallocatechin gallate (ECGC) is a catechin compound from green tea that has been shown to function as a PMEI, putatively binding the PME catalytic site (Lewis *et al*., 2008). Pollen tubes were grown for 3 hours in standard LPGM before replacing the growth media with LPGM supplemented with 0.1mM ECGC. Incubation in LPGM + 0.1mM ECGC for 1 hour did not affect *msl8-5* pollen tube elongation dynamics. WT pollen had fewer pulsatile tubes, from 81.82% in mock to 58.33% in ECGC, although this difference was not significant following Fisher’s exact test (p<0.05). The treated WT tubes that were pulsatile appeared to have greater periods of elongation when treated with ECGC, consistent with the effects of PMEI maintaining extensible esterified homogalacturonan at the tube apex **(Figure 7A, B)**. This ECGC treatment made WT growth dynamics more similar to *msl8*, suggesting that decreased PME activity could induce *msl8*-like phenotypes.

**Figure 7.**
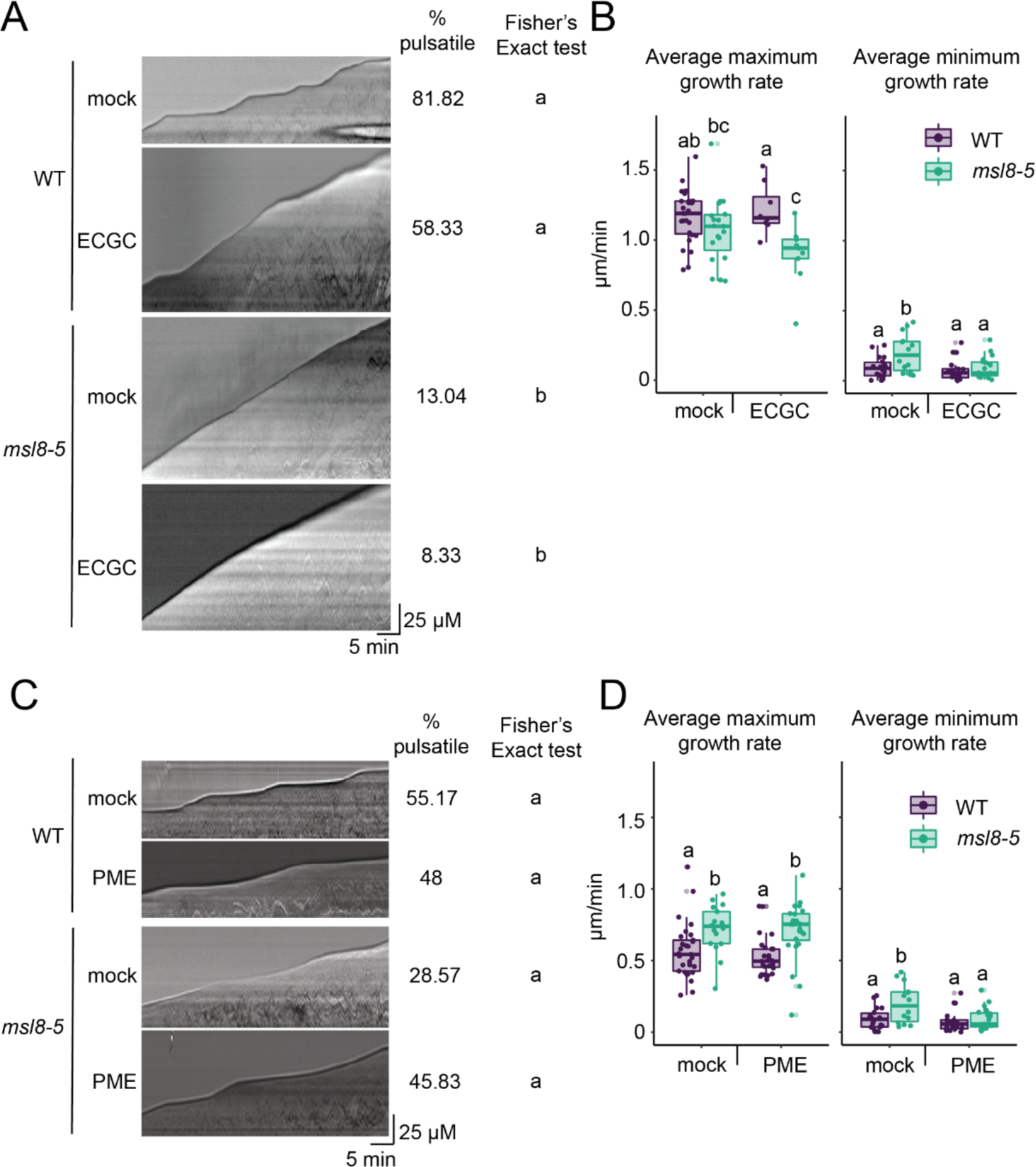
Pectin methylesterase inhibitor treatment causes enhanced elongation in WT, but not *msl8* pollen tubes. **A)** Representative kymographs of pollen tube growth dynamics after one hour treatment with 0.1 mM epigallocatechin gallate (ECGC). Tubes were imaged every 10 sec for 40 min. **B)** Average maximum (left panel) and average minimum (right panel) growth rate from kymograph analysis, n=20-30 tubes per genotype. Statistical analysis by ANOVA followed by Tukey HSD testing and represented by compact letter display, p<0.05. **C)** Representative kymographs of pollen tube growth dynamics in wildtype (top panel) and *msl8-5* (bottom panel) after one hour treatment with 0.1U/µl PME. **D)** Average maximum (left panel) and average minimum (right panel) growth rate from kymograph analysis, n=20-30 tubes per genotype. Statistical analysis by ANOVA followed by Tukey HSD testing and represented by compact letter display, p<0.05.

In the reciprocal test, pollen tubes were grown for 3 hours in standard LPGM before replacing the growth media with LPGM supplemented with 1U/μl orange peel PME. After1 hour in the PME treatment tubes were imaged once every 10 s for 40 min and analyzed by kymograph. The growth rate for both wildtype and *msl8-5* pollen was reduced in mock treatment, indicating a slightly toxic effect from the phosphate buffer the PME was reconstituted in. Orange peel PME treatment did not impact wildtype pollen tubes relative to mock conditions, but it did restore some aspect of pulsatile growth dynamics in *msl8-5* pollen in both mock (28.45%) and treatment (45.83%) conditions, although these changes were not significantly different after a Fisher’s exact test **(Figure 7C, D**). These results indicate that some aspect of the 5mM phosphate buffer (pH 6.5) used in both mock and treatment conditions is both detrimental to pollen tube growth in general and can restore pulsatile growth in msl8 pollen tubes, although it is not clear what the mechanism is behind this effect.

## DISCUSSION

There is substantial correlative data suggesting that the pattern of cell wall deposition and homogalacturonan esterification contributes to pollen tube growth dynamics (Bosch *et al*., 2005; Tian *et al*., 2006; McKenna *et al*., 2009; Winship *et al*., 2010; Rojas *et al*., 2011). Here, we add to these data and introduce the concept of a MS channel influencing these processes by releasing ions to the extracellular space in a membrane tension-dependent manner. Plant MS channels have clearly defined roles in osmoregulation and internal signaling (Basu and Haswell, 2017), but this is the first time that a function for ion flux into the apoplast by plant MS ion channels has been proposed. We found that without MSL8 channel function, pollen tubes lose major growth pauses and have altered homogalacturonan accumulation and esterification at the tube apex. These data support a model whereby MSL8-mediated ion flux modulates pectin extensibility, and thereby elongation dynamics, in the pollen tube apex. This model is described in more detail below.

### Overview of Working Model

We propose that MSL8 is activated in an oscillatory manner during tube elongation. Deformation of the plasma membrane during cell expansion could provide the increase in membrane tension needed to open MSL8 and possibly other MS channels (Dutta and Robinson, 2004; Hepler *et al*., 2013). According to this model, ion flux through MSL8 reduces local cell wall extensibility, which acts as a brake to reduce elongation and initiate a growth pause. The MSL8 braking signal—a membrane tension-induced change in the apoplastic ionome—could promote PME activity or increase pectin crosslinking in order to stiffen the local cell wall.

As mentioned above, it is hypothesized that cell wall deposition or modification rather than turgor fluctuations underlie pulsatile pollen tube growth (Benkert *et al*., 1997; Zerzour *et al*., 2009; Winship *et al*., 2010, 2021; Rojas *et al*., 2011; Van Hemelryck *et al*., 2018; Dumais, 2021). Our results support this hypothesis and position a mechanosensitive ion channel as a possible contributor to oscillatory regulation (**Figure 1**). Notably, pulsatile behavior was not restored in *msl8* pollen tubes provided with osmotic support, indicating that osmotic control does not underlie growth dynamics (**Figure 2**).

### Ion flux through MSL8 is required for its function in oscillatory growth of pollen tubes

MSL8^F720L^ mutant channels, which disrupt ion flux but not channel localization or assembly (Hamilton and Haswell, 2017) do not complement loss of pulsatile dynamics and the altered pectin deposition of *msl8* mutants. (**Figure 1A**, **Figure 3**, **Figure 5A**), suggesting that it is ion flux through the MSL8 pore that is the key function of MSL8 in this context. However, we cannot rule out the alternative explanation that the F720L lesion not only blocks the channel pore, but also prevents it from adopting its open conformation, thereby disrupting a normal interaction with a putative protein signaling partner. While the exact ions passed through MSL8 in the pollen tube apex is not known, it is anion-preferring and Cl^-^ and NO^3-^ are likely candidates.

### Anion efflux as a mechanism of pulsatile growth control

The plant cell wall acts as an ion exchanger between itself and free ions in the apoplastic space, with the wall consisting mainly of negative charges (Shomer *et al*., 2003). For example, H^+^ efflux into the wall is key to the acid growth theory, where acidification of the cell wall results in polymer loosening and extension. However, anions have largely been ignored as potential regulators of cell wall mechanics. While anion efflux at the tube tip appears to be part of the pollen tube growth cycle (Zonia *et al*., 2002), potential roles in the apoplast were not discussed. In one study, chloride treatment of maize leaves resulted in transient alkalinization of the apoplast, which in turn remodeled the apoplastic proteome and stiffened the cell wall (Geilfus *et al*., 2017), suggesting a possible role for anions in cell wall mechanics.

Apoplastic microenvironment chemistry is one area of interest when considering how MSL8-mediated anion efflux might impact pollen tube growth dynamics and pectin status. PMEs are ionically bound to the negatively charged cell wall matrix, and their association and function can be modulated by cation displacement and pH levels (Moustacas *et al*., 1991; Bordenave and Goldberg, 1994; Bosch and Hepler, 2005). Furthermore, PMEI interaction with PMEs has also been shown to be pH sensitive (Bonavita *et al*., 2016). While these studies used tissues and systems other than *A. thaliana* pollen, it is interesting to consider how an anion efflux could locally shift the apoplast electrostatic chemistry in similar but opposing manner, possibly complexing metal cations and reducing their impact on wall bound proteins. Misregulation of PME activity could explain the excess esterified homogalacturonan we observe in tubes lacking MSL8 channel function (**Figure 5A-B**) but does not account for the presence of de-esterified homogalacturonan (**Figure 5C-D**).

### Pectin esterification and crosslinking status as a means of pulsatile growth control

Our observations that tubes lacking MSL8 channel function elongate continuously (**Figure 1A, Supplemental Figure 1**) and have higher levels of esterified homogalacturonan at the tube apex than the WT (**Figure 5A-B, Supplemental Figure 3A**) are consistent with reports that esterified homogalacturonan is more extensible than de-esterified homogalacturonan (McKenna *et al*., 2009; Vogler *et al*., 2015). Together, the observed increase in LM20 signal and the atypical distribution of LM19 signal in *msl8* knock-out and channel-blocked mutant pollen indicate that MSL8 channel function is required for normal pollen tube pectin patterning.

The presence of de-esterified homogalacturonan at the apex of *msl8* mutant pollen tubes could contradict our model. De-esterified homogalacturonan may be present, but not crosslinked into a stiffened matrix that can be detected with our method. Rojas et al (Rojas *et al*., 2011) proposed that chemical relaxation of the wall simultaneously disrupts existing pectic networks while forming new crosslinks with freshly de-esterified polymers as the tube elongates, shifting the load bearing forces to the new pectins until sufficient crosslinking stiffens the pectic gel.

Despite the presence of more de-esterified homogalacturonan at the apex of *msl8* mutant pollen tubes, if it is not sufficiently crosslinked the tube would remain extensible and thus not reach a growth pause, perhaps indicating an imbalance in available Ca^2+^ ions or an overall increase in pectin deposition that outweighs the normal Ca^2+^ pool available.

Our data on PI sensitivity support this, as adding PI inhibits *msl8* tube elongation. PI is proposed to bind the same de-esterified sites as Ca^2+^ on homogalacturonan, but not form crosslinks between polymers (Rounds *et al*., 2011). Thus, increasing PI concentrations can outcompete Ca^2+^ and lead to a destabilized wall that is prone to rupture. Our WT pollen was insensitive to PI treatment, suggesting that there was a sufficient Ca^2+^:de-esterified homogalacturonan ratio that was not disrupted by PI competition. The *msl8* pollen tubes were highly sensitive to PI, showing reduced growth associated with tube tip rupture (**Figure 6**). This indicates an imbalance in Ca^2+^:de-esterified homogalacturonan as mentioned above. One possibility in this scenario is that the excess de-esterified homogalacturonan we detect in *msl8* tubes is already outweighing the amount of Ca^2+^ available to crosslink it, and even a modest amount of PI competition can cause tube rupture.

### Anion efflux might modulate CWI signaling in response to membrane tension and elongation

The CWI pathway monitors and regulates cellular integrity, and in previous work we found a complex genetic relationship between *MSL8* and genes encoding components of the CWI. In germinating pollen grains, MSL8-YFP overexpression could compensate for the bursting defects seen in CWI pathway mutants, and upregulation of CWI signaling could rescue the bursting of *msl8* pollen (Wang *et al*., 2022). The altered cell wall composition at the apex of *msl8* mutant pollen tubes shown here (**Figure 5**), is also a phenotype associated with CWI misregulation (Mecchia *et al*., 2017; Fabrice *et al*., 2018). It is possible that MSL8 anion efflux alters the microenvironment chemistry of the pollen apoplast, thereby impacting interactions between CWI factors such as leucine-rich repeat extension proteins (LRX) and rapid alkalinization factor peptides (RALFs). These interact with each other and with plasma membrane bound receptor proteins to monitor wall status (Ge *et al*., 2017, 2019; Mecchia *et al*., 2017; Fabrice *et al*., 2018) (Moussu *et al*., 2020).

### Future directions

Much remains to be uncovered in defining the mechanism behind MSL8 control of pollen tube growth dynamic and cell wall composition. Dynamic imaging of chloride, pH, and Ca^2+^ fluxes in *msl8* pollen tubes could further define the ionic growth control patterns at play. Defining the specific flux composition of MSL8 channels will be key to formulating testable hypotheses about how MSL8 may be modifying the apoplast microenvironment. Future work could also explore the effect of MSL8 anion efflux on interactions between CWI signaling components, potentially providing an alternative mechanism for regulating cell wall deposition and growth. We feel the results presented here open up a new area of study for the apoplast ionome as a regulatory component of plant cell biology. This positions MS channels as dynamic modulators of the plant cell wall and creates a link translating plasma membrane mechanics to cell wall mechanics. Further exploration of the MSL8-cell wall interaction, as well as the effects of other MS channels in other tissue types is ripe for unlocking greater insights of cell wall dynamics and growth regulation.

## SUPPLEMENTAL FIGURE LEGENDS

**Supplemental Figure 1.** Example kymographs for the indicated lines.

**Supplemental Figure 2.** Quantitative analysis of toluidine blue-stained cell wall patterns. Multiple *msl7* and *msl8* lines (left panel) and multiple of *msl8-5+pMSL8::MSL8* and *msl8-5+pMSL8:: MSL8^F720L^* lines (right panel) are shown. n=90-150 tubes per genotype.

**Supplemental Figure 3.** Extended results of LM20 antibody signal for multiple events of *msl8-5+pMSL8::MSL8* and *msl8-5+pMSL8:: MSL8^F720L^* lines. Some panels repeated from genotypes already shown in Figure 4.

**Supplemental Figure 4.** Extended results of LM20 antibody signal for multiple events of *msl8-5+pMSL8::MSL8* and *msl8-5+pMSL8:: MSL8^F720L^* lines. Some panels repeated from genotypes already shown in Figure 4.

**Supplemental Figure 5.** Cos-488 staining of pectin in live pollen tubes. **A)** Representative images of tubes after 4 hours. Scale bar, 10 μm. **B)** Quantification of signal extending 50 µM from the tip.

## ACKNOWLEDGEMENTS

This work was supported by NSF MCB 1929355. We are especially appreciative of helpful and insightful conversations with Dr. Daniel Damineli on the technical implementation of the CHUKNORRIS software as well as discussion of oscillatory dynamics and quantifications. We additionally thank Michael Dyer and the Jeanette Goldfarb Plant Growth Facility at Washington University for assistance with plant growth.

